# Epiviz File Server: Query, Transform and Interactively Explore Data from Indexed Genomic Files

**DOI:** 10.1101/865295

**Authors:** Jayaram Kancherla, Yifan Yang, Hyeyun Chae, Hector Corrada Bravo

## Abstract

Genomic data repositories like The Cancer Genome Atlas (TCGA), Encyclopedia of DNA Elements (ENCODE), Bioconductor’s AnnotationHub and ExperimentHub etc., provide public access to large amounts of genomic data as flat files. Researchers often download a subset of files data from these repositories to perform their data analysis. As these data repositories become larger, researchers often face bottlenecks in their exploratory data analysis. Based on the concepts of a NoDB paradigm, we developed epivizFileServer, a Python library that implements an in-situ data query system for local or remotely hosted indexed genomic files, not only for visualization but also data manipulation. The File Server library decouples data from analysis workflows and provides an abstract interface to define computations independent of the location, format or structure of the file.

## 1 Introduction

Genomic data repositories like The Cancer Genome Atlas (The Cancer Genome Atlas Research Network et al. 2013), Encyclopedia of DNA Elements (Davis et al. 2018) (ENCODE), Bioconductor’s (Huber et al. 2015) AnnotationHub (Morgan et al. 2019) and ExperimentHub (bioconductor 2019) etc., provide public access to large amounts of genomic data as flat files. Researchers often download a subset of files data from these repositories to perform their data analysis. As these data repositories become larger, researchers often face bottlenecks in their exploratory data analysis. Increasing data size requires longer time to download, pre-process and load files into a database, to run queries efficiently.

Interactive visualization of data can be a powerful tool to enable exploratory analysis. As users get familiar with the data and gain insights, it would be even more efficient to interactively hypothesize, validate, visualize and compute the intermediate results of the analysis. Currently available interactive visualization tools for genomic data, namely genome browsers, fall into two broad categories. One that uses a database management system to load genomic data from files into tables, create indices or partitions for faster query of data by genomic intervals. The other category of genome browsers query data directly from indexed genomic file formats like BigBed, BigWig (Kent et al. 2010) or Tabix (Li 2011). However these tools are limited only to exploration of data from files.

Based on the concepts of a NoDB paradigm (Alagiannis et al. 2012), we developed epivizFileServer, a Python library that implements an in-situ data query system for local or remotely hosted indexed genomic files, not only for visualization but also data manipulation. Our design of the File Server library was based on the following goals

1. Efficiently parse minimal necessary bytes from an indexed genomic file to query data for a specific genomic region
2. Define transformations and summarizations directly over files and lazily compute these at query time
3. Scale operations to concurrently process multiple file query and transformation requests
4. Implement cache over files for faster access and improve repeat query performance
5. REST API for developers and bioinformaticians to build interactive visualization and exploratory tools over genomic data stored in flat files
6. Integration with existing bioinformatic tools and software to interactively visualize and explore genomic data directly from files

The File Server library decouples data from analysis workflows and provides an abstract interface to define computations independent of the location, format or structure of the file. Our major contribution on this research project was to efficiently and intuitively define transformations and summarizations directly over files, without the hassle of downloading the files locally or preprocess for exploratory data analysis.

Using the library, researchers and analysts can author shareable and reproducible data exploration workflows in an intuitive and programmatic way. Using the library, users can query and explore data directly from local and publicly hosted indexed genomic files. If the files are hosted on a public server, the library requires the server hosting the data files to support HTTP range requests (Lafon, Fielding, and Reschke 2014), so that the parser can only request the minimum necessary byte-ranges needed to process the query. The library supports various file formats - BigBed, BigWig and any tabular file that can be indexed using Tabix. Once these data files are described, users can define summarizations and transformations on these files using NumPy (or NumPy-like) functions.

The File Server library uses Dask (*Dask: Scalable analytics in Python*) to scale and concurrently process multiple query and transformations over files. The cache implementation makes sure we only access bytes not already accessed and stored locally. Developers of bioinformatic tools and systems can use the Library’s REST API to build interactive data visualization or exploratory tools over files. We demonstrate the integration in the use cases section of this paper by integration with Galaxy (Afgan et al. 2018), a widely used open source bioinformatic web platform for analysis of genomic data and Epiviz (Chelaru et al. 2014), an interactive and integrative functional genome browser to visualize and explore these datasets. The browser supports various charts to explore genomic data, heatmap and scatter plots to visualize gene expression, block (linear and stacked) tracks for visualizing genomic regions of interest and line tracks (stacked, multi stacked) for visualizing signal (ChIP-seq, methylation etc) data. Hovering over a region in one visualization highlights this region in other tracks providing instant visual feedback to the user. These visualizations are developed using standards based web component framework, are highly customizable, reusable and can be integrated with most frameworks that support HTML (Kancherla et al. 2018). Figure 1 describes a high level overview of these components in the Epiviz File Server library.

**Figure 1:**
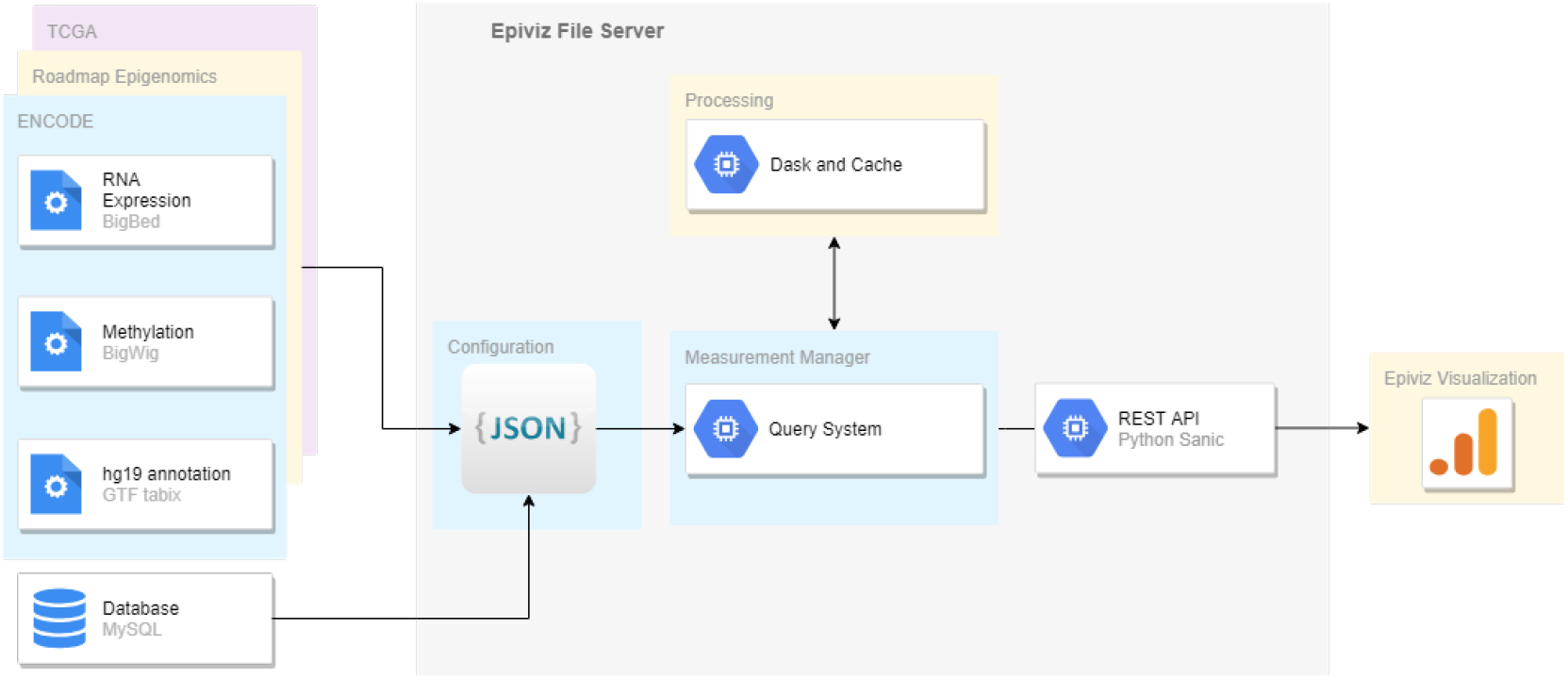
A high level overview of the Epiviz File Server Library. Several publicly hosted genomic data repositories provide access to various experiments as files. Epiviz File Server library supports directly querying these indexed genomic files. The files can be imported into the system either manually or using a JSON configuration file for large datasets. Datasets are described in the measurements module and provides a programmatic interface to query and parse files. This module allows users to define transformations over files using any NumPy function and lazily compute these transformations at run time. We use Dask to scale and manage the queries and computations over files. The cache layer makes sure we only request for bytes not already accessed and reduces the number of file access requests. The entire system can also be accessed using a REST API. Developers can use the API to build interactive visualization and exploration tools. One such example is Epiviz, an interactive statistical visualization and exploration system for functional genomics data.

## 2 Methods

### 2.1 Naive Approach

A naive approach to implement a query system over genomic files would be to first import the files into a database management system like MySQL. Accessing data from a database provides various advantages, we can query the data quickly by indexing on intervals (genomic position), a standard declarative query language like SQL, and being able to modify the table schema on-the-fly. As the size of the data increases, there is a significant cost to initialize, load and prepare the dataset for queries. Our experience importing a histone modification data file from a ChIP-seq experiment into a database significantly increased the data-to-query time i.e., time to import the file into a table, time to index the table, and time to query the table for a particular genomic region. Even after this, the database table had to be custom configured to use advanced indexing schemes like partitions and index by partition to improve the query performance. This is not a feasible approach to individually tune and optimize query performance when dealing with large public repositories like the NIH roadmap epigenomics (Bernstein et al. 2010) or ENCODE that host datasets from hundreds of experiments.

Assuming we do have all the data files imported into a database system, performing transformations on data directly is limited to the functions implemented in the system. Also if we need to perform complex summarization or transformations across genomic datasets, we would need to first perform interval overlap operations that are limited or non-existent in database systems. To work around these limitations, one would implement backend programming logic (middleware) to first query the database for multiple datasets, perform interval overlap operations to align the data by genomic location, summarize and then apply transformations.

### 2.2 (our) NoDB Approach

NoDB is a new database design philosophy to make database systems more accessible and reduce the data-to-query time (Alagiannis et al. 2012). In the naive approach, we expect the data-to-query time to significantly increase as the size of the data increases. The NoDB philosophy is a way of querying the data files directly, in-situ, hence removing the loading time. A simple approach would be to load the data file into memory every time we need to query for a particular region. This is not efficient for 1) repeated query processing since we would load the same file into memory on every request and 2) memory usage especially when querying several large data files at the same time. In order to work around the issue of loading the entire file into memory, we need to create and store an index based on genomic positions, query this index and only access the minimum necessary bytes of the file to process the query.

The genomics community has long used specialized binary file formats like bigBed, bigWig or Tabix for quickly accessing particular regions of the genome. These file formats contain an index which allows us to quickly and efficiently access blocks of the file that contains the data, then only parse these blocks to process the query. These formats also support remote file access, allowing multiple parallel requests to process at the same time. Our NoDB approach uses these indexed genomic file formats as the base for in-situ query processing. Some of these file formats also summarize the data at different zoom levels (base-pair resolution) which is extremely useful especially for interactive exploration and visualization.

#### 2.2.1 Related Work

Several existing genome browsers and tools support visualization of genomic data from flat files. These include the UCSC Genome Browser (Kent et al. 2002), Dalliance (Down, Piipari, and Hubbard 2011), JBrowse (Buels et al. 2016), Integrated Genome Browser (Freese, Norris, and Loraine 2016) etc. However, these tools only visualize genomic data from files and do not perform transformations over data. Integrated Genome Viewer (Robinson et al. 2011) allows users to combine tracks through its interface and perform Add, Subtract, Multiply or Divide operations but are tightly coupled to the visualization.

In addition to querying data from files, Epiviz File Server provides an interface to compute transformations over data directly from files. The library decouples data analysis from visualization workflows. This way new transformations can be applied over files and can be explored interactively using Epiviz.

#### 2.2.2 Traditional Database server vs file-based web server

To efficiently process queries, a traditional database server implements cache management and can provide scalability as the system demands. A file based web server always has to access the files to query for a genomic region. This may not be efficient for repeated query processing for the same region. When querying BigWig and BigBed using the tools (bigWigSummary and bigBedSummary) from UCSC utilities (*UCSC Utilities*), it implements a url cache layer using the sparse file feature available in UNIX based systems, which downloads and create files locally as blocks of the file are accessed. This may not be efficient when accessing hundreds of files from a repository.

Epiviz File Server library implements a cache management system to store frequently accessed blocks as part of the file objects. This reduces the number of requests to the file (if available locally) or server hosting the file. As queries are processed, we store the binary blocks of the file that were recently and frequently accessed. The cache allows the library to quickly process repeated queries for a genomic region. As the number of queries and files increase, it also increases the cache size and impacts memory usage. The library implements task schedulers that automatically serialize (pickle) file objects from memory to disk. This process efficiently manages memory and frees up system resources for other tasks. After storing a file object to disk, if a new request is made to query for a genomic region from the same file, we load (deserialize) the file object into memory and process the query.

For scalability, our library uses Dask, a lightweight distributed computing library for Python. Dask manages, distributes and schedules tasks dynamically. The Dask scheduler is asynchronous and event driven, simultaneously responding to requests for computation from multiple clients. This provides flexibility to concurrently handle a variety of workloads from multiple users at the same time while also handling node failures and additions. In addition, the File Server library uses Sanic (*Sanic* 2019), an asynchronous library to make API queries to files and integrates well with the Dask system for handling web requests.

#### 2.2.3 Transformations over files (Computed Measurements)

As previously stated, if an analyst needs to perform complex summarization or transformations across genomic datasets, they would need to first perform interval overlap operations that are limited or non-existent in database systems. Transformations over data are also limited in scope to the methods implemented in database management systems. Existing genome browsers that support file based visualization are only limited to exploration of data. As an illustrative example, if a researcher is exploring ChIP-seq data for a particular histone (*H3k24me3*) marker across different tissues, and would like to visualize the difference in binding across these tissues, the typical way is to use a computational environment or tools to read the files, align the data from these files to apply transformations, compute the difference and then store the dataset as a file or into a database. For the purpose of exploration, it would be very efficient to be able to define these transformations over files without having to precompute and interactively explore these transformations.

The Epiviz File Server library allows computing transformations over files using methods available in NumPy or custom functions using a NumPy compatible definition (called computed measurements). The transformations are not pre-computed but lazily evaluated at query time. When querying these computed measurements for a particular genomic region, we first query the individual files for data, perform interval overlap operation to align the datasets and then apply the transformation. We use Dask to scale and compute transformations. The code documentation and the use case sections provides more details on defining transformations over files.

#### 2.2.4 Optimizations

The file server library performs a number of optimizations mostly when querying for data from a file that contains zoom levels (BigWig and BigBed). This is extremely useful especially for interactive visualization and exploration. For example, visualizing signal data from a roadmap ChIP-Seq dataset for a 10000 base-pair (bp) region, on a webpage canvas of size 800 (width) by 400 (height) pixels. if the visualization library plots one bp per pixel, it would be very inefficient to try to render 10000 data points in an 800 pixel width screen. In such scenarios, the library automatically chooses the appropriate zoom level to query for data. If zoom levels are not available, the library automatically summarizes the data into smaller bins and computes a mean across each bin. This decreases the response to query time, improves rendering performance and is efficient for interactive visualization and data exploration for large genomic datasets.

#### 2.2.5 Data Import

When accessing files from genomic data repositories that contain hundreds of files from various experiments, it is inefficient to individually describe these files. We implemented several import functions to batch load data files from local or public repositories.

To access files available through Bioconductor’s AnnotationHub or ExperimentHub, we implemented methods to directly import these into the library.

The UCSC Genome Browser provides Track hubs (Raney et al. 2014), a useful tool for visualizing large number of genome wide data sets. Track hubs are web-accessible directories that contain genomic data and can be viewed on the Genome Browser. Methods are available to load track hub repositories into the Epiviz File Server library.

Epiviz File Server library also supports a JSON based configuration library to define files. The configuration file defines a collection of files and for each file, describes the location of the file (public or full local path), its format, name, and annotations. An example configuration file is described in the use case section (Figure 2).

**Figure 2:**
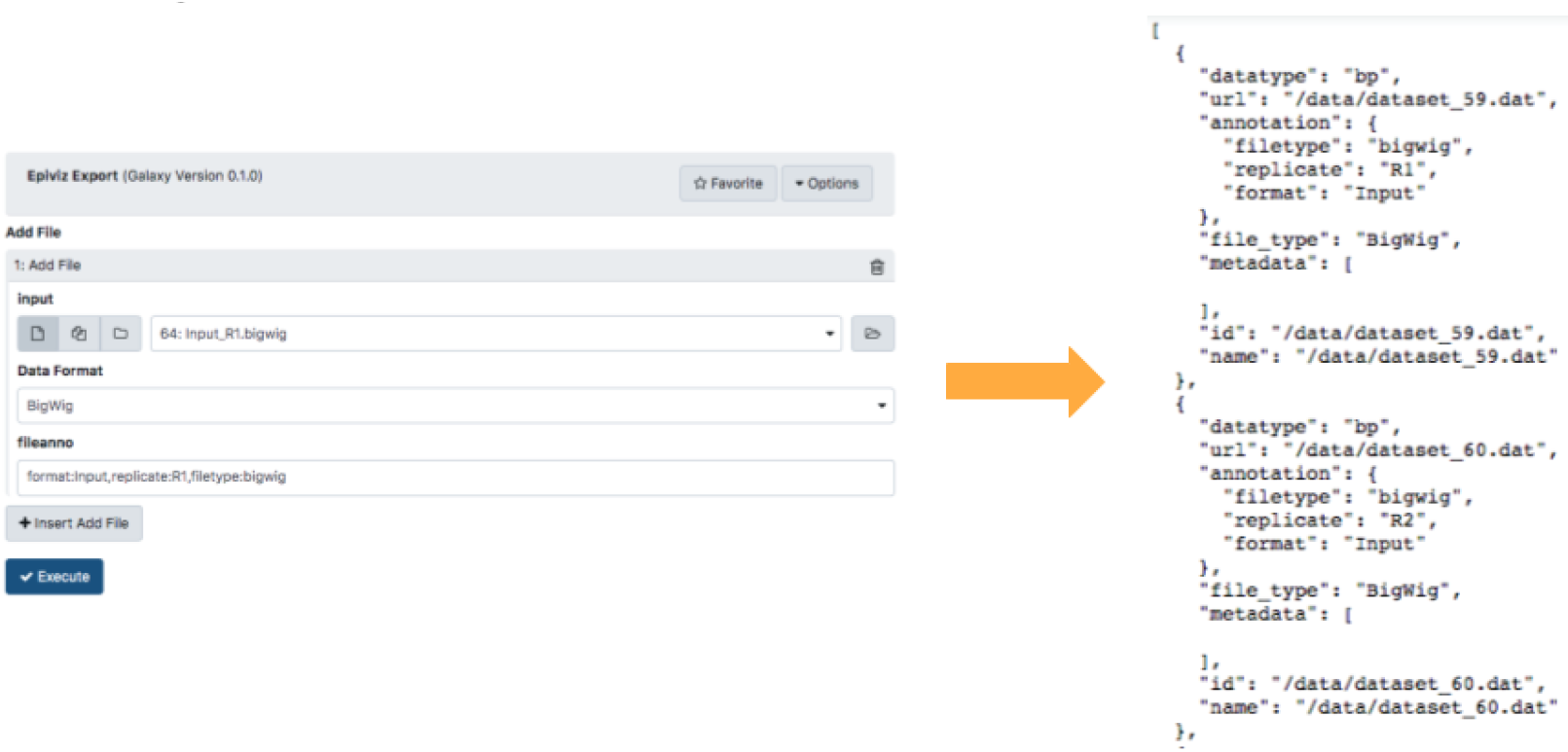
Screenshot of the Epiviz Galaxy Tool for defining files. On the left is a screenshot of the tool to select multiple files generated from the galaxy workflows. Options are available to set annotations and format for each file. On the right, a configuration file is generated for epiviz file server to import these files.

Regardless of how these files are defined and imported, users can always define transformations over the files.

#### 2.2.6 File Formats

Epiviz file server currently supports indexed genomic file formats - BigWig, BigBed, bam (with bai), sam (with sai) or any tabular data file that can be indexed using Tabix. These files either have an index as part of the same file (BigBed, BigWig etc) or create a separate index file (Tabix). The index (usually a BTree or an RTree) allows the library to quickly navigate (or seek) and only access the blocks of the file that contains the data for a genomic region. These formats also support remote file access, allowing multiple parallel requests to process at the same time.

## 3 Use cases

### 3.1 Integration with Galaxy workflows

Galaxy is one of the widely used open source web-based platforms for analysis of genomic data. Galaxy aims to make computational biology accessible to researchers with less programming experience. It has an easy to use user interface to create reproducible and shareable workflows and installs various bioinformatic tools without all the complexity. At its core, Galaxy is a file based workflow platform, every step in the workflow creates a file(s) and these files are used as inputs in the next step. Our goal with integrating epiviz with galaxy is to create a single computational environment where users can analyze and explore datasets generated by Galaxy workflows. Galaxy provides Galaxy Interactive Environments, a framework to integrate external tools with Galaxy workflows and user interface.

To integrate Epiviz with Galaxy, we need to 1) Register and Run Epiviz as a tool with the Galaxy system 2) Define and access files generated at various steps in a galaxy workflow and 3) Query interactively visualize the files using Epiviz.

#### 3.1.1 Register and Run Epiviz with Galaxy

To integrate external tools, Galaxy provides a framework called Galaxy Interactive Environments (IE) (GrÃŒning et al. 2017). The first step in IE is to create a docker container for epiviz to manage its dependencies internally and also efficiently manage system resources. The docker container hosts both the epiviz file server (the library to manage and query files) and the epiviz user interface to interactively query and visualize datasets. Once the container is configured, IE requires a configuration file (mako) that spins up the docker instance on demand and run epiviz inside galaxy. The mako configuration file sets various parameters for the docker image and configures the ports to use from the image to serve the user interface. The file server library also needs access to the files generated during the galaxy workflow. Instead of copying over files to the docker instance, we mount the data directory used by galaxy to the docker image.

#### 3.1.2 Define files from galaxy workflows

To define and access files to visualize using Epiviz, we created a Galaxy Tool (Blankenberg et al. 2014) that the user integrates with their final step of the workflow. This tool allows the user to choose various files generated in their workflow, define annotations and file formats, and generates an Epiviz configuration file as shown in Figure 2. Epiviz File Server library running in the docker image from the previous step uses this configuration file to load these files into the instance.

#### 3.1.3 Visualize files in Galaxy workflow

After generating the configuration file in the previous step, users can visualize these datasets using epiviz within the galaxy interface. This displays the epiviz application running inside the docker container on Galaxy and users can now visualize and interactively explore various files. Figure 3 illustrates the process of integration and describes various steps in the process.

**Figure 3:**
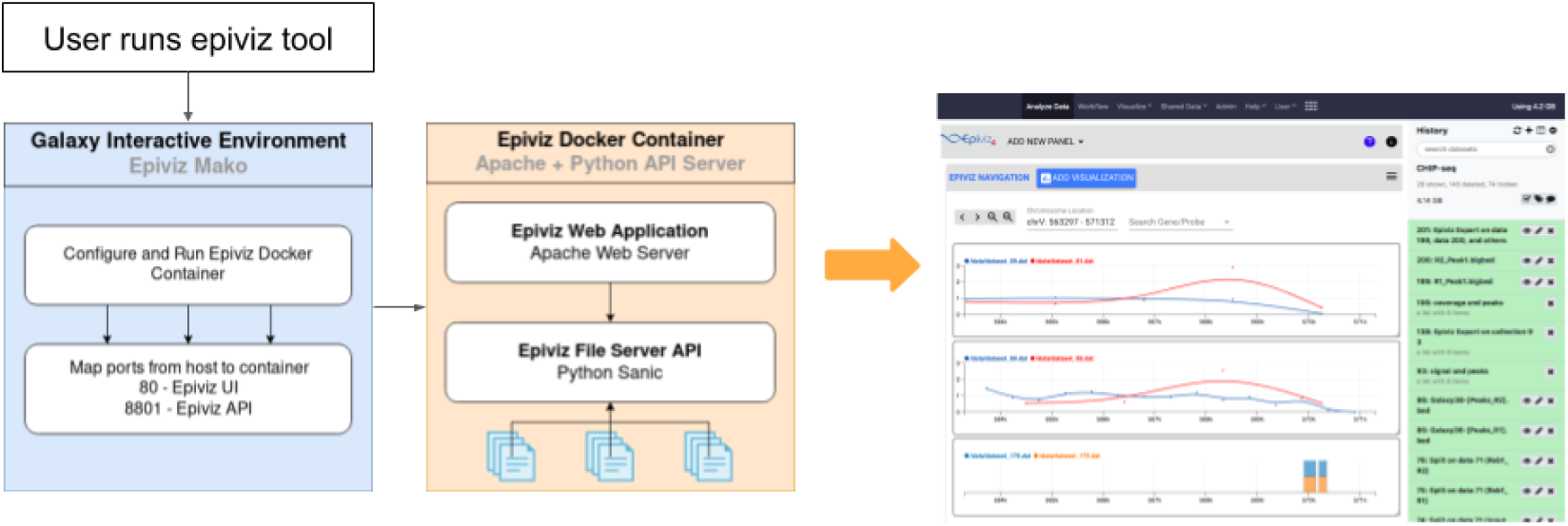
Overview of Epiviz integration with Galaxy. After running a workflow, users use the Epiviz Galaxy Tool to choose files and define annotations to generate an epiviz configuration file. When the user chooses epiviz to visualize this file, Galaxy understands that epiviz is a Galaxy Interactive Environment (IE). Galaxy using the configuration defined in the IE (mako file), spins the epiviz docker instance. The epiviz docker instance runs both the file server library and the epiviz user interface to interactively visualize and explore genomic files. Once the docker image loads, Galaxy now serves the user interface from the instance on its User Interface as shown on the right. Users can now interactively visualize and explore various files chosen in the epiviz tool (first step in this process).

To demonstrate the integration, we use the Analysis of ChIP-seq data workflow from Galaxy (*Analysis of ChIP-seq data*). The workflow uses data where immunoprecipitation was performed with antibodies from *Reb1. Reb1* recognizes a specific sequence (*TTACCCG*) and is involved in many aspects of transcriptional regulation by all three yeast RNA polymerases and promotes formation of nucleosome-free regions (NFRs) (Hartley and Madhani 2009; Raisner et al. 2005). After executing the entire workflow, users can now run the Epiviz tool to generate the configuration file on the coverage and peak calling files. Figure 4 shows the datasets visualized from the results of this workflow.

**Figure 4:**
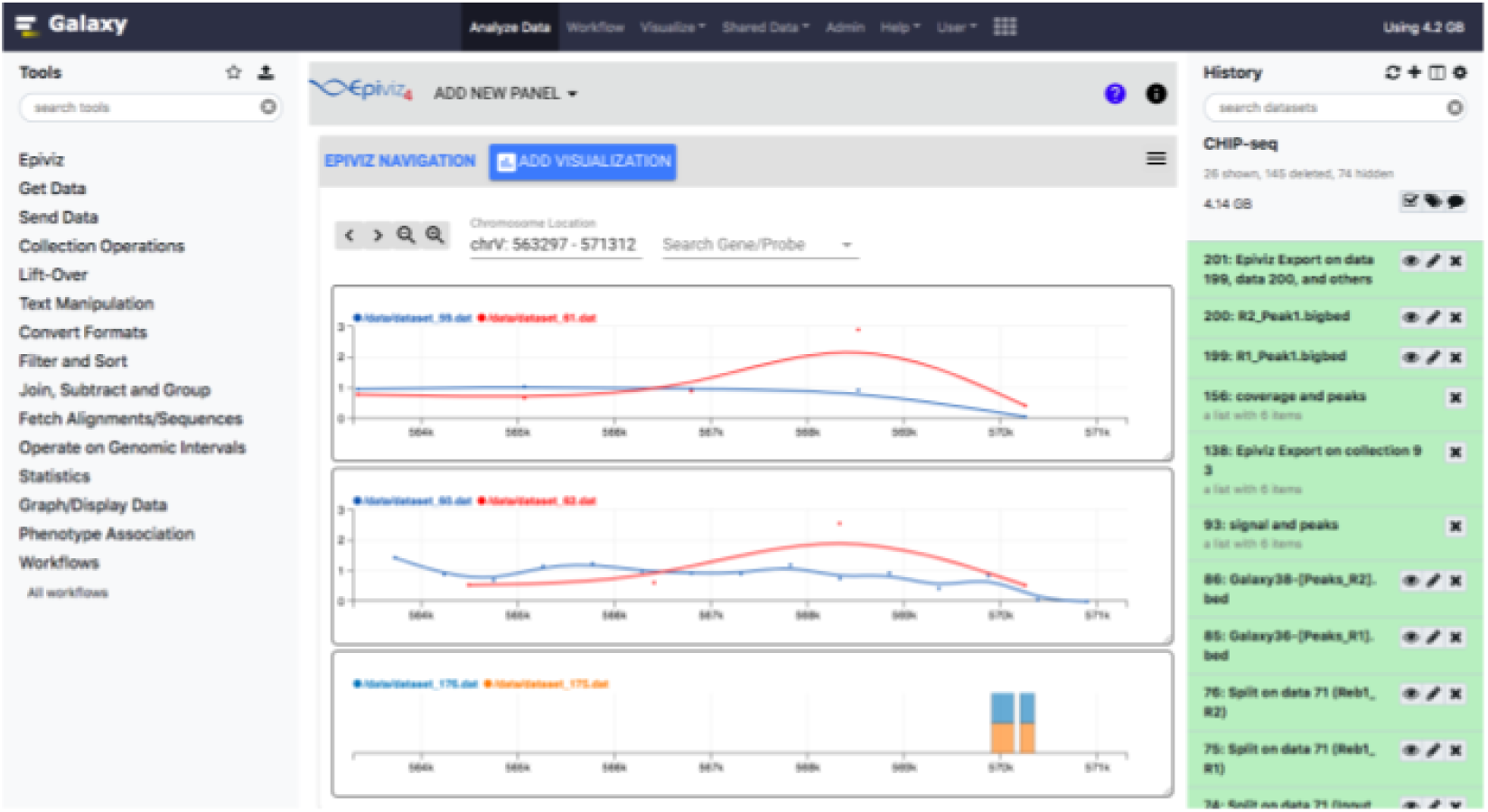
Epiviz visualizing signal and peak data from the ChIP-seq galaxy workflow. (top to bottom) The first track visualizes Replicate 1 and its Input signal, the second track visualizes Replicate 2 and its Input signal. After running the MACS2 peak calling tool on the dataset, the Galaxy workflow generates peak files and the last track visualizes these peak regions across both the replicates.

### 3.2 Roadmap

The NIH Roadmap Epigenomics Mapping Consortium leverages next-generation sequencing technologies to map DNA methylation, histone modifications, chromatin accessibility and small RNA transcripts in tissues, selected to represent the normal counterparts of tissues and organ systems frequently involved in human disease. The data files from this consortium are available at http://egg2.wustl.edu/roadmap/web_portal/ or http://www.roadmapepigenomics.org/.

For this use case, we use BioConductor’s AnnotationHub to query for data files that are part of the NIH Roadmap Epigenomics project. The API for AnnotationHub is hosted at http://annotationhub.biocon-ductor.org/. We downloaded the AnnotationHub sqlite database to extract information for all available resources in the hub. We then query this database for resources associated with the roadmap project. We get a total of 9932 data resources from AnnotationHub. These include DNA methylation signal, ChIP seq fold change signal and pvalues for various tissues and histone markers. To easily import these AnnotationHub records into the fileserver, we added a helper function ‘import_ahub’. We filtered samples for the brain region and defined transformations over these files. We define difference in histone modification between different brain tissues for this dataset, for example, difference in *H3K27me3* histone modification across “*Brain Hippocampus Middle*” and “*Brain Inferior Temporal Lobe*”. We setup an instance of epiviz interactive visualization tool (http://54.82.116.84/browser/) on Amazon Web Services (AWS) to explore the roadmap dataset and the transformations as shown in Figure 5.

**Figure 5:**
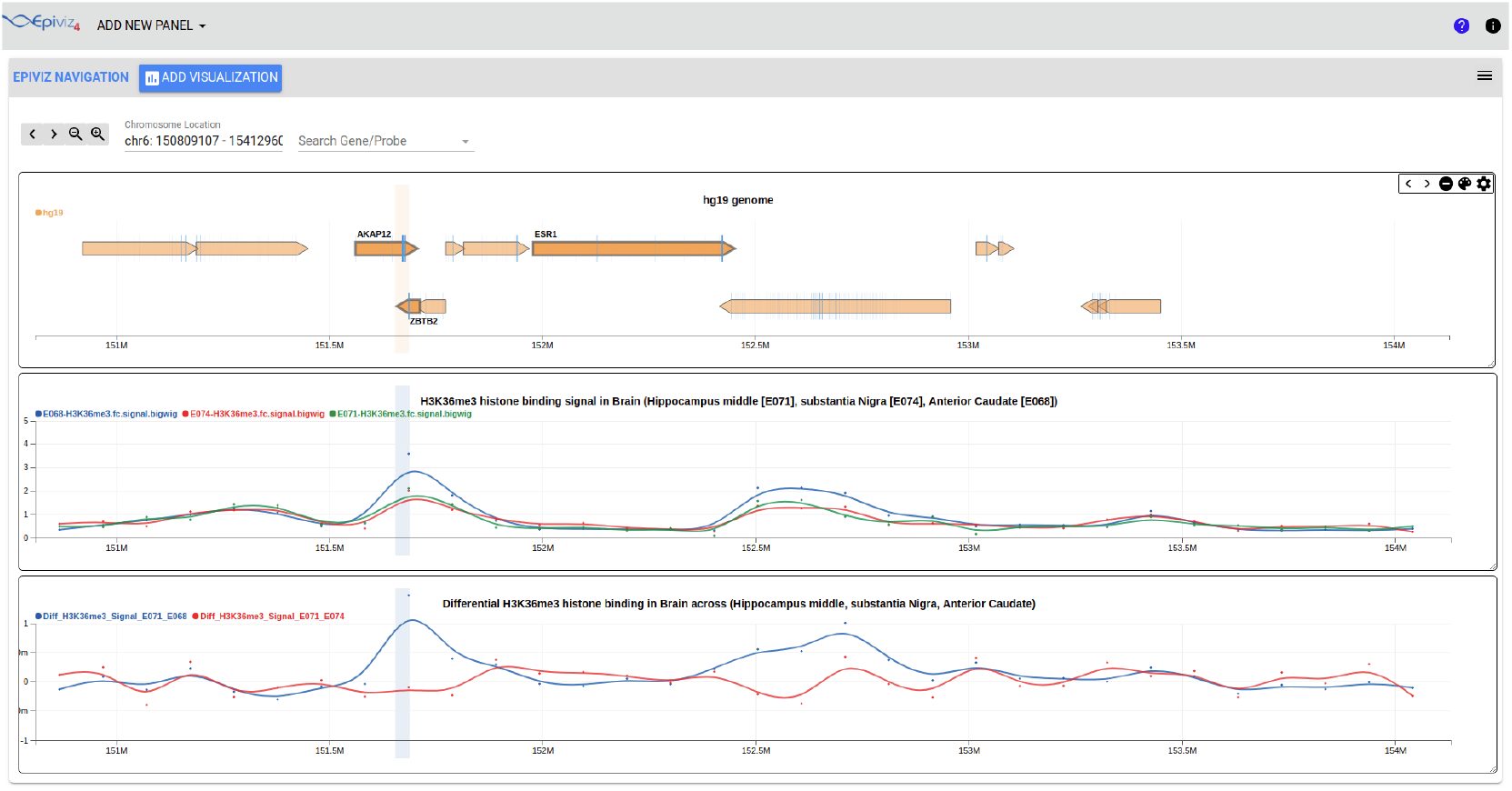
Interactive visualization of data from the NIH Roadmap Epigenomics project. This figure demonstrates the Epiviz File Server library querying and computing transformations over data available from the NIH Roadmap Epigenomics project. We chose the *ESR1* and its neighboring gene region for this example. (top to bottom) The first track is a genome annotation track visualizing various genes in the given genomic region for the *hg19* human genome. The line track in the middle is visualizing the *H3K36me3* binding signal from the ChIP-seq experiments across three different brain tissues (*Brain Hippocampus Middle* [E071], *Brain Substantia Nigra* [E074], *Brain Anterior Caudate* [E068]). This track queries the data directly from the files. The last line track is a transformation over files to compute difference in histone binding across different tissues. Computations can be defined over files and are lazily evaluated at run time. The last track is visualizing these computations, the difference in *H3K26me3* binding between different brain tissues. This workspace is available at http://54.82.116.84/browser/

## 4 Software Availability

Epiviz File Server is open source and the source code is available on GitHub at http://github.com/epiviz/epivizFileParser

The documentation for the File Server library is available at http://epivizfileparser.rtfd.io

The package is published to PyPI and is available at http://pypi.org/project/epivizFileServer

For integration with galaxy, we have two different GitHub repositories, one for the Epiviz Tool (http://github.com/epiviz/epivizGalaxyTool) and the other for Epiviz Interactive Environment (http://github.com/epiviz/epivizGalaxyIE). The repositories also contain instructions to setup the Interactive environments in Galaxy.

For the use case describing the Roadmap dataset, the code is available in the File Server GitHub repository inside the use cases folder. The AWS instance is accessible at http://54.82.116.84/browser/

The code for benchmarks is also available in the same repository inside the benchmarks folder.

The code is available under MIT License.

## 5 Benchmarks

We performed several tests to evaluate 1) Performance with and without the cache implementation and 2) Overhead in lazily evaluating transformations.

All tests were run on a standard Amazon AWS EC2 (t2.xlarge) instance with 4 vCPUs and 16GB memory.

### 5.1 Impact of Cache on performance

To evaluate the impact of cache on the system, we randomly generated 20 different genomic range queries and repeatedly queried these against the web server for 60 seconds. We use wrk (Glozer 2019), a HTTP benchmarking tool capable of generating significant load to test the API. We run the tool on its default settings using 2 threads and 10 connections concurrently to send requests to the system.

We run the system on two different modes, one with the cache feature and the other without the cache. In the cache implementation, if the given genomic region already exists in the cache, it is fetched quickly and sent back to the user whereas in the non-cache setting, the library always parses the file to query data for the given genomic range. Since local file access is fast, our results are comparable between the cache and non-cache settings. Instead, we hosted the files on a S3 object store bucket at University of Maryland, adding network latency to the system. The results indicate the cache implementation significantly improves the performance of the system. Table 1 displays the results of these tests. To make sure other processes are not interfering in the benchmarking process, we disabled the serialization process for file objects discussed in the methods section.

**Table 1:** Impact of cache on processing requests. This table displays displays the Average Latency and the Requests processed per second measured when benchmarking the File Server API with and without a cache implementation. The tests were run using the wrk tool by repeatedly querying 20 different genomic range queries for 60 seconds. With cache, the library was able to process a significantly higher number of genomic range queries resulting in higher throughput and lower latency.

| Implementation | Avg Latency (in ms) (± SD) | Requests (per sec) |
| --- | --- | --- |
| No Cache | 1440 (± 430.43) | 2.00 |
| Cache | 104.52 (± 196.81) | 132 |

### 5.2 Overhead in lazily computing transformations

Epiviz File Server library lazily computes transformations or summarizations directly over files. We measure the overhead in computing transformations run time as opposed to pre-computing a transformation, storing and querying this file. For this test, we choose 20 different genomic datasets (bigwig files) from roadmap, we created an instance of the file server that at runtime computes a mean signal value with increasing number of files starting from 2 and upto 20 genomic datasets. In addition, we also precomputed the mean signal using WiggleTools (Zerbino et al. 2014) and store these files. This allows us to compare the overhead in run-time computation versus the pre-compute.

We run similar benchmarks as before using the wrk tool, where we randomly generate 5 different genomic range queries and query the system for 60 seconds (2 threads and 10 connections). We measure the Average Latency and Requests processed per second by the system across five different runs. We calculate the mean and standard deviation for the measured metrics. Our results are shown in Figure 6. As expected, as the transformation involves more files, the latency of the system increases hence serving fewer requests per second compared to directly querying a pre-computed file. However, the system is still interactive with reasonable query response times.

**Figure 6:**
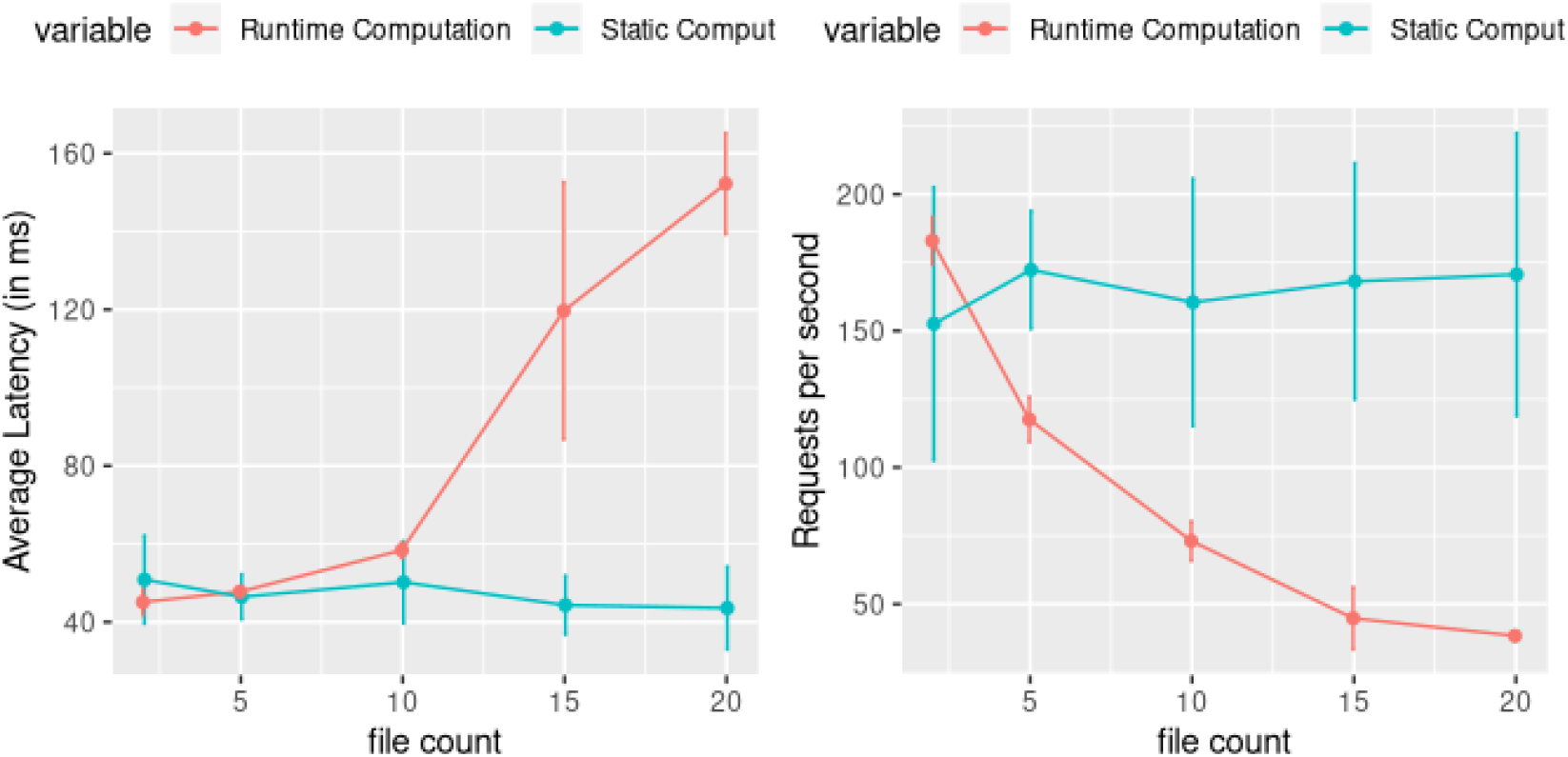
Impact of computing transformations at run time. Since the Epiviz File Server library computes transformations at run time, we measure the overhead in computing transformations lazily versus querying a file that stores the pre-computed result. We ran the wrk tool to benchmark the system on 5 genomic range queries for 60 seconds using 2 threads and 10 connections. We measure the Average Latency and Requests processed per second by the system across five different runs. We also calculate the mean and standard deviation for the measured metrics. The results indicate the latency of the system increases as we increase the number of files involved in the computation, hence lowering the number of requests processed per second. However, the system is still interactive with reasonable query response times.

## 6 Discussion

Epiviz File Server is a Python library to interactively query and transform data directly from indexed genomic files. The library decouples data from analysis workflows and provides an abstract interface to define computations independent of the location, format or structure of the file.

Genomic files hosted on public repositories are fairly stable and often do not change. Also when the library queries the file for data, it first parses the index of these files. Given these conditions, we can either pre-merge the index of all files or use a data structure to dynamically update index as we parse files. This would be similar to the interval data structure used by GIGGLE (Layer et al. 2018), which uses a B+ tree to create an index from thousands of genomic data and annotation files. We would like to extend the File Server to use giggle or giggle like structures for querying data, or an index structure that is updated dynamically as more queries are processed across files.

Several interval data structures in genomics are optimized for quickly querying data for a given genomic region. But new technologies like Single cell experiments generate large datasets measuring tens of thousands of features over thousands of single cells. Although very efficient to query by genomic region, if we are only interested in a few cells from such large matrices, the library still has to parse the entire row and filter the columns. The Tabix format is a commonly used indexing technique for any tabular genomic data set (the first three columns must be chromosome location, start and end). This includes RNA-seq data (csv), SAM or BAM files etc. Interactive analysis, including visualization of these datasets is a challenging task especially for queries to filter by columns (or cells) and efficiently transferring these long matrices between server and client. Piccolo et al. 2019 recommend using coordinate based fixed width formats as a fast and scalable approach to query tabular genomic data. In addition, the genomics community has been using HDF5 based formats AnnData or H5ad (Wolf, Angerer, and Theis 2018) and loom (Loom) to store large genomics datasets. We are currently exploring ways to efficiently query HDF5, H5ad or loom format files. These formats do not natively support remote querying like BigWig or Tabix but require an additional server like h5serv (Heber 2013) setup for web queries.

Our group also developed Metaviz (Wagner et al. 2018), an interactive and statistical data exploration tool for metagenomic datasets. Instead of the linear organization of data usually found in genomic datasets, features of metagenomic datasets are mapped hierarchically to a taxonomic reference database. This also changes the linear navigation implemented in genome browsers to adapt to a hierarchical based navigation of taxonomic features. This work also extends to single cell datasets, where the hierarchy can be defined based on the similarity of cells using methods like Clustering Trees (Zappia and Oshlack 2018). We are currently exploring ways to use HDF5 based formats like loompy (*loompy* 2019) to include this hierarchical structure within the file and be able to query and explore the data.

The File Server library uses Pandas to perform interval overlap operations on genomic data and for compatibility with libraries like Dask to scale transformations over data. When computing across different datasets, we would first perform interval overlap operations and then apply the transformation. Recent approaches like Pyranges (Stovner and SÃŠtrom 2019) have implemented data structures for efficiently manipulating genomic intervals in Python.

We would extend the library to incorporate these file formats and data structures to efficiently query and interactively explore multi-omic datasets.

## 7 Conclusion

Based on the concepts of a NoDB paradigm, we present a file based Python library, an in-situ data query system for indexed genomic files, not only for visualization but also transformation. The library implements several features provided by a traditional database system for query, transformation, visualization and caching. The use cases demonstrate the flexibility in integrating the File Server library with existing bioinformatic tools and public repositories. We discussed new research approaches to build a comprehensive file based data visualization and exploration system for genomics datasets.

